# Hyperglycemia enhances cancer immune evasion by inducing alternative macrophage polarization through increased O-GlcNAcylation

**DOI:** 10.1101/831610

**Authors:** Natália Rodrigues Mantuano, Michal A. Stanczak, Isadora de Araújo Oliveira, Nicole Kirchhammer, Alessandra Filardy, Gianni Monaco, Ronan Christian Santos, Agatha Carlos Fonseca, Miguel Fontes, César de Souza Bastos, Wagner B. Dias, Alfred Zippelius, Adriane R. Todeschini, Heinz Läubli

## Abstract

Diabetes mellitus (DM) significantly increases the risk for cancer and cancer progression. Hyperglycemia is the defining characteristic of DM and tightly correlates with a poor prognosis in cancer patients. The hexosamine biosynthetic pathway (HBP) is emerging as a pivotal cascade linking high glucose, tumor progression and impaired immune function. Here we show that enhanced glucose flow through the HBP drives cancer progression and immune evasion by increasing O-GlcNAcylation in tumor-associated macrophages (TAMs). Increased O-GlcNAc skewed macrophage polarization to a M2-like phenotype. HBP or O-GlcNAcylation inhibition reprogrammed TAMs to an anti-tumoral phenotype. Finally, we found an upregulation of M2 markers on TAMs in DM2 patients with colorectal cancer compared to non-diabetic normoglycemic patients. Our results provide evidence for a new and targetable mechanism of cancer immune evasion in patients with hyperglycemia, advocating for strict control of hyperglycemia in cancer patients.

**Significance:** Hyperglycemia increases O-GlcNAc levels in TAMs, programing them to a pro-tumorigenic phenotype (M2-like), contributing to cancer progression. Inhibition of O-GlcNAcylation could therefore be used to reprogram intratumoral macrophages to an anti-tumoral phenotype.

## Introduction

Diabetes mellitus (DM) is one of the most common metabolic disorders with a high morbidity and mortality leading to uncontrolled hyperglycemia [1]. Prevalence of DM is increasing at an alarming rate and on a global scale [2]. Epidemiological evidence shows that DM patients display a significantly higher risk of developing multiple types of cancers [3–6] and also demonstrated that cancer recurrence, metastasis or fatal outcome occurs with much higher incidence in cancer patients with hyperglycemia [7–9]. Although an association of a poor prognosis with hyperglycemia in cancer patients has been shown, mechanistic studies often focus on metabolic changes in tumor cells [10]. Studies on the influence of hyperglycemia on the tumor immune microenvironment are often lacking and further investigations are needed.

Glucose metabolism, and in particular the hexosamine biosynthetic pathway (HBP), emerged as an important cascade linking cancer progression and hyperglycemia [10–13]. The HBP provides UDP-N-acetyl-D-glucosamine (UDP-GlcNAc) and its derivatives for glycosylation using glucose, glutamine, Acetyl-CoA and UTP as substrates [14]. UDP-GlcNAc is highly responsive to cell nutrient fluctuations as its synthesis depends on products of the several metabolic pathways and is therefore considered to be a metabolic sensor [14]. Our recent work showed that increasing extracellular glucose concentrations provoke aberrant glycosylation, increasing cell proliferation and invasion *in vivo* due to increased flux through the HBP [14]. Hyperglycemia increased levels of intracellular O-GlcNAcylation in tumor cells [12], a post-translational modification of intracellular proteins catalyzed by O-GlcNAc transferase [15] [15] and counteracted by O-GlcNAcase (OGA) [16]. Increase of O-GlcNAc levels induced by high glucose can affect the expression of genes and deregulation of biochemical pathways to respond to environmental changes [10].

Metabolic changes can strongly influence immune cell function [17–21]. While glucose is needed for proper effector function as T cell activation and M1 macrophage polarization [21, 22], it is largely unknown how a systemic increase of glucose in circulation of diabetic cancer patients instructs the local tumor immune microenvironment. In order to improve our understanding of the influence of hyperglycemia, we used a mouse model of hyperglycemia to study the changes in the tumor immune microenvironment. We characterized alterations of cell infiltration, activation, and polarization and provide a molecular mechanism for the effect of hyperglycemia on tumor growth.

## Materials and Methods

### Study approval

All experiments involving mice were approved by the local ethical committees in Basel, Switzerland (approval number 2747) and in Rio de Janeiro, Brazil (approval number IBCCF214-09/16) and performed in accordance with the regulations. All human tumor samples were anonymized and obtained from the Marcílio Dias Hospital following informed consent (approval 39063014.8.0000.5256, Brazil).

### Cell culture

Mouse colon adenocarcinoma MC38 cells were donated by Dr. M. Smyth (QIMR Biomedical Research Institute, Brisbane, Australia) and mouse melanoma B16D5 cells by Dr. L. Weiner (Georgetown University, USA). Tumor cell lines were cultured in Dulbecco’s Modified Eagle Medium (DMEM, 25 mM Glucose, Sigma-Aldrich), supplemented with 2 mM L-glutamine, 1 mM sodium pyruvate, 1% MEM non-essential amino acids, 500 U/ml streptomycin/penicillin and 10% heat-inactivated fetal bovine serum (Sigma-Aldrich) at 37°C and 5% CO_2_. All cells were tested for mycoplasma by PCR.

### Mice

Male C57BL/6 and NOD scid gamma (NSG) mice were obtained from Janvier Labs [23], bred in-house at the Department of Biomedicine, University Hospital Basel, Switzerland and were used at 10–12 weeks old. Hyperglycemia (HyG) was induced with a single intraperitoneal (i.p.) injection of 150 mg/kg streptozotocin (STZ, Sigma-Aldrich) diluted with 0.1 mM sodium citrate buffer as vehicle (pH 4.3). Euglycemic control mice (EuG) were treated with vehicle only. Seven days after STZ injection, 5×10^5^ syngeneic tumor cells in 100 µl phosphate-buffered saline (PBS) were injected subcutaneously into the flank of mice. Tumor growth and health scores were measured 3 times per week until a maximum size of 1500 mm^3^ (advanced tumor stage), or 300–400 mm^3^ (early tumor stage). Perpendicular tumor diameters were measured using a caliper, and tumor volume was calculated according to the following formula: tumor volume (mm^3^) = (d^2^*D)/2, where d and D are the shortest and longest diameters (in mm) of the tumor, respectively. Mice that developed ulcerated tumors were sacrificed and excluded from further analysis.

### Blood glucose measurement

Blood sampling was performed from the tail vein seven days after STZ injection and on day of tumor resection. Glucose levels were measured using a validated glucose meter (FreeStyle lite, Abbott). STZ-treated mice were considered to be HyG if the blood glucose levels exceeded 250 mg/dl at the end of the experiment.

### Intratumoral glucose measurement

Tumors were collected, weighed, cut in small pieces and mechanically dissociated using a micropipette. Each tumor was resuspended in 1 ml PBS, filtered, glucose concentrations measured by the glucose colorimetric detection Kit (Invitrogen) and normalized to tumor mass.

### Treatments

Depletion of intra-tumoral macrophages was achieved by i.p. anti-CSFR1 antibody treatment (CD115, clone AFS98, BioXCell) with a total of 7 injections, 1 mg before tumor inoculation, followed by 6 injections of 400 µg each every 3 to 4 days over 3 weeks.

HBP inhibition *in vivo* was achieved by treatment of tumor bearing mice with 3 doses (10 mg/kg) of DON (6-Diazo-5-oxo-L-norleucine, Sigma-Aldrich) every 3 days, beginning when tumors reached 300–400 mm^3^. In addition, mice were treated with a single i.p. injection of DON (10 mg/kg) or the OGA-inhibitor Thiamet G (TMG, 20 mg/kg, Sigma-Aldrich) and animals sacrificed 2 days after the treatment.

### Tumor digestion

Tumors were surgically collected, cut in small pieces and digested in a solution of collagenase IV (Worthington), DNase I (Sigma-Aldrich), Hyaluronidase (Sigma-Aldrich) and accutase (PAA Laboratories) for 1 hour at 37°C. After incubation, cells were filtered, washed, separated by density centrifugation with Histopaque-1119 (Sigma-Aldrich) and cell suspensions frozen at −80°C for future analyses. All samples were stained using LIVE/DEAD Fixable Blue Dye (Invitrogen) and various panels of antibodies to identify different population of immune cells. All samples were acquired with the Fortessa LSR II Flow Cytometer (BD Biosciences) and analyzed in the FlowJo software (BD Biosciences).

### Bone marrow derived macrophages (BMDM) and treatments

BMDMs were differentiated in petri dishes with M-CSF (20 ng/ml, Peprotech) or 20% L929 supernatant for 6 days in high glucose (HG, 25 mM) or low glucose (LG, 5mM) DMEM. Cells were plated at 5×10^5^/well in 12 well-plates and treated for 24 hours with 10 µM of the inhibitors DON, TMG or OSMI-1 (Sigma-Aldrich). BMDMs were polarized to M1 with IFN-γ 50 ng/ml (Peprotech) and LPS 20 ng/ml (Sigma-Aldrich) or to M2 with IL-4 20 ng/ml (Peprotech) for 48 hours. BMDMs were washed in FACS buffer, followed by incubation with anti-CD16/CD32 blocking antibodies (eBioscience) for Fc-receptor blocking. Polarization was confirmed by staining against CD206, CD86 and MHC-II. Cells were also collected for metabolite extraction and immunoblotting.

### Metabolite extraction and nucleotide sugar analysis

Polar metabolites were extracted from 5×10^6^ macrophages with chloroform, methanol and water (1:1:1). Each sample was subjected to chromatographic separation on a Hypercarb PGC column (ThermoFisher) running in HPLC Prominence (Shimadzu). Nucleotide sugars were eluted using a discontinuous linear gradient of mobile phases A (0.2% formic acid and 0.75% ammonium hydroxide in water) and B (95% acetonitrile with 0.1% formic acid and 0.07% ammonium hydroxide). The detection was achieved using a diode array detector (SPM-M20A Prominence, Shimadzu) set to 250 to 400 nm, coupled to a Q-TOF mass spectrometer (maXis Impact, Bruker Daltonics). The retention times of nucleotide sugars were previously established using standards. The chromatograms and mass spectra were analyzed with DataAnalysis 4.2 (Bruker Daltonics). UDP-GlcNAc concentration was quantified by measuring the area of its correspondent UV peak at 262 nm normalized to the total area of the chromatogram from 5 to 30 min, using a calibration curve. For the other nucleotide sugars, the peaks of their respective extracted ion chromatograms were evaluated similarly.

### Immunoblotting

BMDMs were washed with PBS and homogenized in lysis buffer (150 mM NaCl, 30 mM, Tris-HCl, pH 7.6, 1 mM EDTA, 1 mM EGTA, 0.1% SDS, 1 mM phenylmethylsulfonyl fluoride, and 1 μM O-(2-acetamido-2-deoxy-d-glucopyranosylidene)amino-N-phenylcarbamate (PUGNAc). Cell lysates were sonicated and centrifuged. Supernatant was collected, protein concentration was determined and modified Laemmli buffer was added. Samples were separated on SDS-polyacrylamide gels and were subsequently electroblotted to nitrocellulose membrane (Bio-Rad). The membranes were blocked in Tris-buffered saline with 0.1% (v/v) Tween 20 with either 3% (w/v) bovine serum albumin or 3% (w/v) nonfat dry milk. The blocked membranes were then incubated overnight at 4 °C with primary antibodies against O-GlcNAc (CTD 110.6 or RL-2, Santa Cruz), arginase-1 (clone H52, Santa Cruz), NOS2 (clone N20, Santa Cruz) and β-actin (clone 13E5, Cell Signaling). The blots were then washed, incubated with the appropriate secondary antibody, developed using ECL (GE Healthcare) and exposed to Image Quant LAS 4000 (GE Healthcare). ImageJ software was used for densitometric analysis of immunoblots and measurements were normalized against β-actin.

### Histology and immunofluorescence

Paraffin sections of CRC patients with diabetes mellitus and control patients were obtained from the Marine Hospital Marcílio Dias (HNMD), Rio de Janeiro, Brazil. All samples were from surgical resections and were evaluated by a board-certified pathologist. Sections were deparaffinized automatically and antigen retrieval was performed in citrate buffer. Tissues were permeabilized with 0.1% Triton X-100 and blocked with 1% BSA. Primary antibodies were incubated overnight in 1% BSA solution. The antibodies used were anti-CD68 (clone Y1-82A, BD Biosciences), anti-CD206 (polyclonal, Abcam, Cat.: 64693) and anti-CD163 (polyclonal, Abcam, Cat.: 87099). After incubation, slides were washed and incubated with the respective secondary antibody and then washed. Slides were further stained DAPI (ThermoFisher) and mounted with ProLong diamond antifade mountant. Pictures were taken in a Nikon Eclipse Ti-S microscope and analyzed using ImageJ.

### Statistics

Graphs and statistical tests were done with Prism 8 (GraphPad). Statistical significance between 2 groups was tested using paired Student’s t tests. Comparison between multiple time points were analyzed by one-way ANOVAs with multiple comparisons and Tukey post-test. Differences in survival curves were analyzed by the Gehan-Breslow-Wilcoxon test.

## Results

### Hyperglycemia increases tumor growth and reduces survival in two different syngeneic mouse models

In order to study the influence of systemic hyperglycemia on tumor progression, we used a mouse model of acute hyperglycemia by applying the cytotoxic drug streptozotocin (STZ), destroying pancreatic beta cells [24]. We tested the growth of MC38 and highly metastatic mouse melanoma B16D5 cells subcutaneously transplanted into C57BL/6 mice treated with STZ (Fig. 1A). Survival was significantly shorter and tumor growth increased in hyperglycemic mice bearing subcutaneous B16D5 tumors (Fig. 1B, C). Similarly, hyperglycemic mice injected with MC38 tumor cells showed a significantly shorter survival than euglycemic mice (Fig. 1D, E). In addition, hyperglycemia was observed in the peripheral blood as well as within the extracellular space of the tumor microenvironment, demonstrated by increased glucose concentrations in the interstitial tumor fluid (Fig. 1F). These experiments demonstrate that hyperglycemia increases tumor growth and thereby reduces survival in two different syngeneic mouse models.

**Figure 1:**
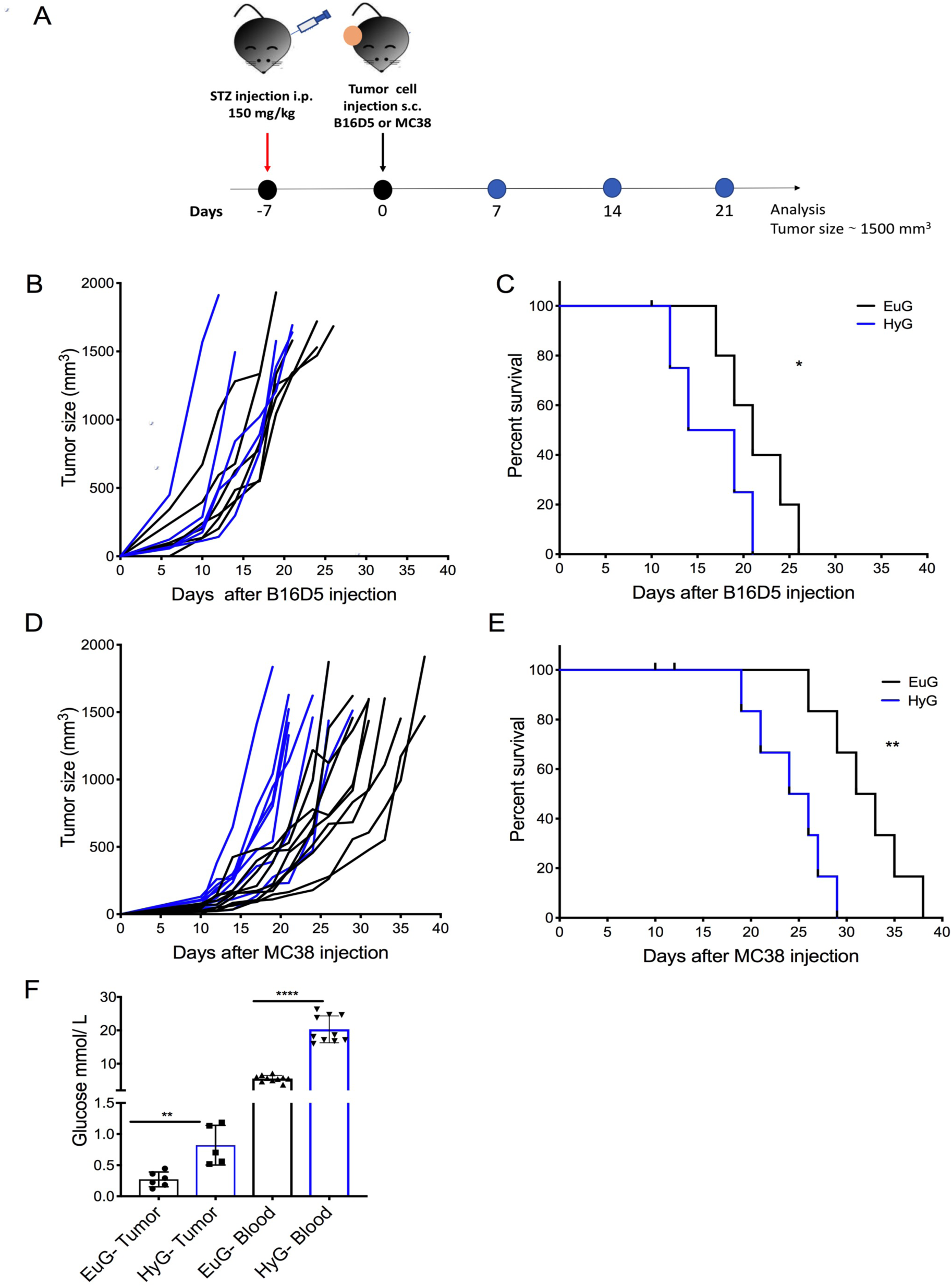
Hyperglycemia increases tumor growth. Hyperglycemia (HyG) was induced *in vivo* in C57BL/6 mice with an i.p. injection of 150 mg/kg STZ. (A) Euglyemic control mice (EuG) received vehicle in a single i.p. injection. Seven days after STZ injection, MC38 or B16D5 cancer cells were injected subcutaneously. Tumor growth was measured 3 times per week until reach ∼1500 mm^3^. (B) Growth curves and (C) Survival curves of B16D5 (n=6 EuG and 5 HyG) and (D, F) MC38 tumors (n=10 EuG and 10 HyG). p values were calculated using the Gehan-Breslow-Wilcoxon test *P < 0.05 and ** P < 0.01. (F) Glucose was measured in the tumor microenvironment of MC38 tumors (n=6 EuG and 5 HyG). Statistical analyses were performed by paired Student’s t test **P < 0.01.

### Sialylation is not a key mechanism in hyperglycemia-induced tumor growth

Recent studies have shown that hyperglycemia can lead to an increase of sialylation in tumor microenvironment [12]. More importantly, hypersialylation of tumors can mediate immune evasion by engaging the sialoglycan-Siglec immune checkpoint [25, 26]. In our model systems, hyperglycemia induced an increase in sialylation demonstrated by increased Siglec-E-Fc binding when compared with cells grown in normoglycemia (Fig. S1A). We therefore tested the role of sialylation in the hyperglycemia model by using GlcNAc-epimerase (GNE) deficient MC38 tumor cells, which display a strong reduction in cell surface sialylation [25]. As previously published [25], GNE-deficient MC38 cells grew slower compared to wild type cells. Yet, hyperglycemia still increased tumor growth, conserving the key difference in tumor growth between hyperglycemic and euglycemic mice (Fig. S1B). This finding was confirmed in mice deficient for Siglec-E, which showed similar tumor growth as wild-type animals, but were smaller in the hyperglycemic compared to euglycemic mice (Fig. S1C). We concluded that engagement of the sialoglycan-Siglec pathway is not the main mechanism of enhanced tumor growth in hyperglycemic mice.

### Hyperglycemia skews intratumoral macrophage polarization

To delineate the cellular immune mechanisms involved in hyperglycemia-induced tumor progression, we characterized the immune cell infiltrates in size-matched tumors by flow cytometry (see gating strategy, Fig. S2). We found a significant reduction of CD8^+^ tumor-infiltrating lymphocytes (TILs, Fig. 2A), as well as of conventional CD11c^+^MHC-II^+^ dendritic cells (Fig. S3A) in tumors from hyperglycemic mice. We found a reduction of CD4^+^CD25^+^FOXP3^+^ regulatory T cells (Fig. 2B), while no significant differences in the frequencies of CD4^+^ TILs (Fig. 2C) and B cells (Fig. 2D) and were observed in the infiltrates of hyperglycemic tumors. In addition, we found that the total percentages of CD3^+^ cells were the same among both groups (Fig. S3B). The presence of Ly6G^+^ myeloid-derived suppressor cells (MDSC) was decreased in tumors from hyperglycemic mice (Fig. S3C), while no difference was seen in the Ly6C^+^ MDSCs (Fig. S3D). The numbers of CD11b^+^F4/80^+^Gr1^−^ tumor-associated macrophages (TAMs) was similar in both conditions (Fig. S3E). However, a significant increase in CD206 expression was found in hyperglycemic MC38 (Fig. 2E, S3F) and B16D5 tumors (Fig. S3I). This result was accompanied by a decrease of the M1 marker MHC-II in MC38 (Fig. 2F, S3G), but not B16D5 tumors (Fig. S3J), while MHC-II^+^CD206^+^ cells were not altered in both cases (Fig. S3H, S3K).

**Figure 2:**
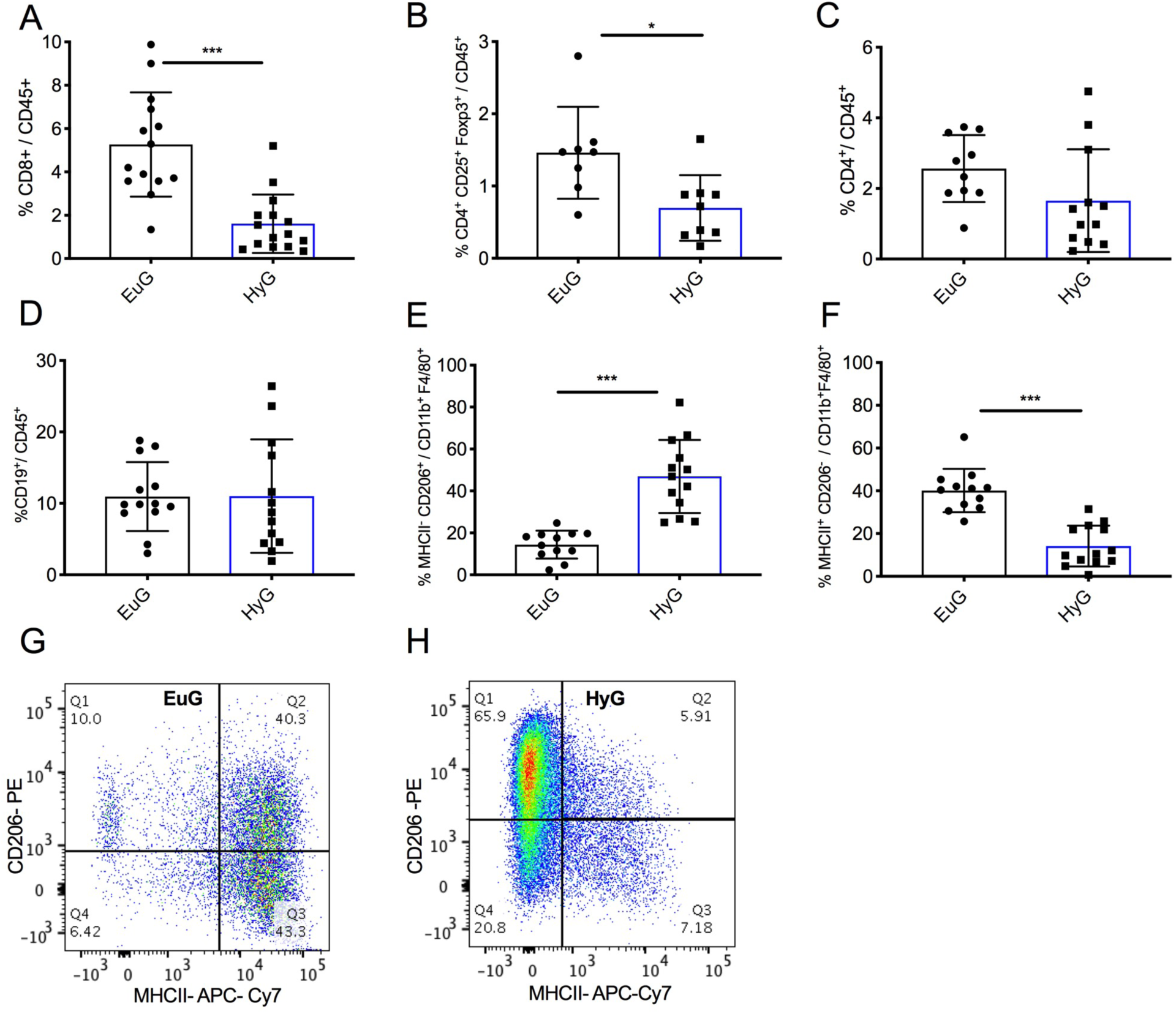
Hyperglycemia skews intratumoral macrophage polarization. Hyperglycemia (HyG) was induced in *vivo* by intraperitoneal injection of streptozotocin (STZ) 150 mg/kg, while Euglyemic control (EuG) received the vehicle. Seven days after STZ treatment, MC38 cancer cells were injected subcutaneously. Tumor growth was measured 3 times per week until the late time point and the inflammatory infiltrated cells were analyzed by flow cytometry. Tumor infiltrated lymphocytes (TILs) were analyzed and previously gated on CD3^+^ cells (n=10–12), (A) CD8^+^ TILs, (B) Regulatory T cells CD4^+^CD25^+^FoxP3^+^ and (C) CD4+ cells were performed on CD45^+^ cells (n=7–9). (D) The percentage of tumor infiltrated B cells infiltrated, gated on CD19^+^ cells (n=13). Tumor associated macrophages (TAM) were gated on CD45^+^F4/80^+^CD11b^+^Gr1^−^ cells and polarization was accessed by MHC-II and CD206 labeling (E, F). Representative dot plot from (G) EuG (n=12) and (H) HyG (n=13) mice. Results are expressed as mean ± S.D. p values were calculated using the Students’ t-test. * P < 0.05; *** P < 0.001.

Taken together, the analysis of inflammatory infiltrates suggests that hyperglycemia impacts the immune response against the tumor by changing the tumor immune microenvironment.

### Tumor associated macrophages mediate immune evasion in hyperglycemic mice

Next, we aimed to determine the functional contribution of the immune system and specific immune cell types to the observed effect of hyperglycemia-promoted tumor growth. Therefore, we induced hyperglycemia in NOD scid gamma (NSG) mice that are devoid of a functional immune system. Tumor growth of subcutaneously injected MC38 cells (Fig. 3A) was faster than in immunocompetent mice (Fig. 3B), while no differences were observed between hyperglycemic and euglycemic mice (Figure 3A). This finding indicates that growth difference observed in our mouse model were mainly mediated by changes in the immune response. Further characterization by gene expression analysis showed a decreased presence of inflammation-associated genes, using immunocompetent C57bl/6 mice. Genes involved in immune cell recruitment and activation such as *Il1b*, *Cxcl1*, *Ccl5*, *S100a9* and *S100a8* were significantly downregulated in hyperglycemic tumors compared to euglycemic tumors (Fig. S4A). In contrast, genes involved in tumor-promoting inflammation as the M2 macrophage marker *Cd163* and *Cd276*, an immune checkpoint member, were increased supporting an important function of macrophages in this model (Fig. S4A). We therefore tested the contribution of macrophages to the hyperglycemia-induced tumor growth by treating mice with anti-CSF1R antibodies. The treatment of mice with anti-CSF1R showed a significant reduction of intratumoral macrophages (Fig. 3C–E). Hyperglycemic mice treated with anti-CSF1R showed similar tumor growth as treated euglycemic mice. The effect of hyperglycemia was abrogated by anti-CSF1R treatment and the resulting reduction in intra-tumoral macrophages. These results suggest a key role for intumoral macrophages in mediating the effect of hyperglycemia on tumor growth and suggest an interplay between innate and adaptive responses in the observed effect.

**Figure 3:**
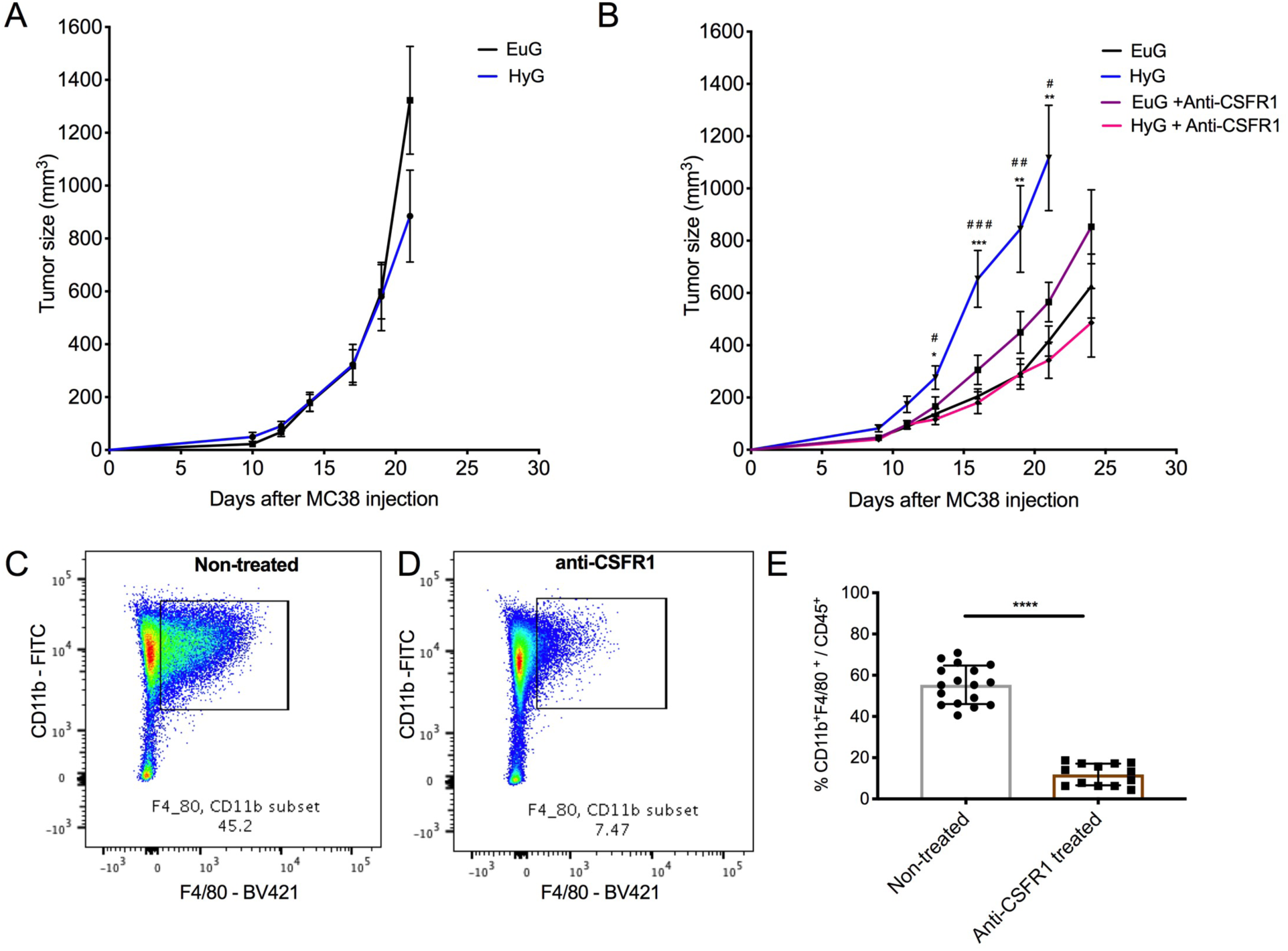
Tumor associated macrophages mediate immune evasion in hyperglycemic mice. Hyperglycemia (HyG) was induced *in vivo* in C57BL/6 mice with an i.p. injection of 150 mg/kg STZ. Euglyemic control mice (EuG) received vehicle in a single i.p. injection. Seven days after the STZ injection, (A) MC38 cancer cells were implanted subcutaneously in NOD SCID mice and tumor area was measured. (B) The macrophage reduction was carried out by injection of anti-CSFR1 antibody in HyG (n=6–10) and EuG (n=7–12) C57BL/6 mice. In total 7 injections, 1 mg before MC38 injection and 6 more injections (400 g) each 3 or 4 days. #. p values were calculated using Anova one-way with Tukey post-test. * *P* < 0.05, ** *P* < 0.01, *** *P* < 0.001, where * HyG vs. EuG and # HyG vs. HyG + Anti-CSFR1. Representative dot plots of macrophage reduction comparing (C) non-treated mice and (D) CSFR1 treated. (E) The frequency of CD11b^+^F4/80^+^ inside the CD45^+^ population (n=13–18). Results are expressed as mean ± S.D. p values were calculated using the Students’ t-test ****P<0.0001.

### Flux through the hexosamine biosynthetic pathway induces M2 polarization

Hyperglycemia can increase the rate of flux through the HBP, resulting in increased levels of UDP-GlcNAc in cells in high glucose environments [12]. Previous work has shown that UDP-GlcNAc is a characteristic of M2 polarization [28] and acts as a substrate for O-GlcNAcylation [14]. In addition, increased O-GlcNAcylation has been demonstrated to influence macrophage function [29–31]. We therefore investigated whether hyperglycemia could change macrophage polarization through increased O-GlcNAcylation. First, we tested whether polarization of *in vitro* cultured BMDMs would be affected changed by the glucose concentration of the cell culture medium (Fig. S4B). High glucose medium augmented UDP-GlcNAc levels (Fig. 4A) and O-GlcNAcylation in BMDMs, with the highest levels being reached under M2-polarizing conditions (Fig. 4B, S4C). Other activated monosaccharides were differentially affected by high glucose, with CMP-Neu5Ac levels decreasing under high glucose conditions (Fig. S4D-F). High glucose changed the macrophage phenotype measured as a decrease of the M1 markers CD86 (Fig. 4C, F, I), MHC-II (Fig. 4D, G, J), and NOS2 (Fig. S4G), while M2 markers such as CD206 (Fig. 4E, H, K) and Arginase1 (Fig. S4G) were increased.

**Figure 4:**
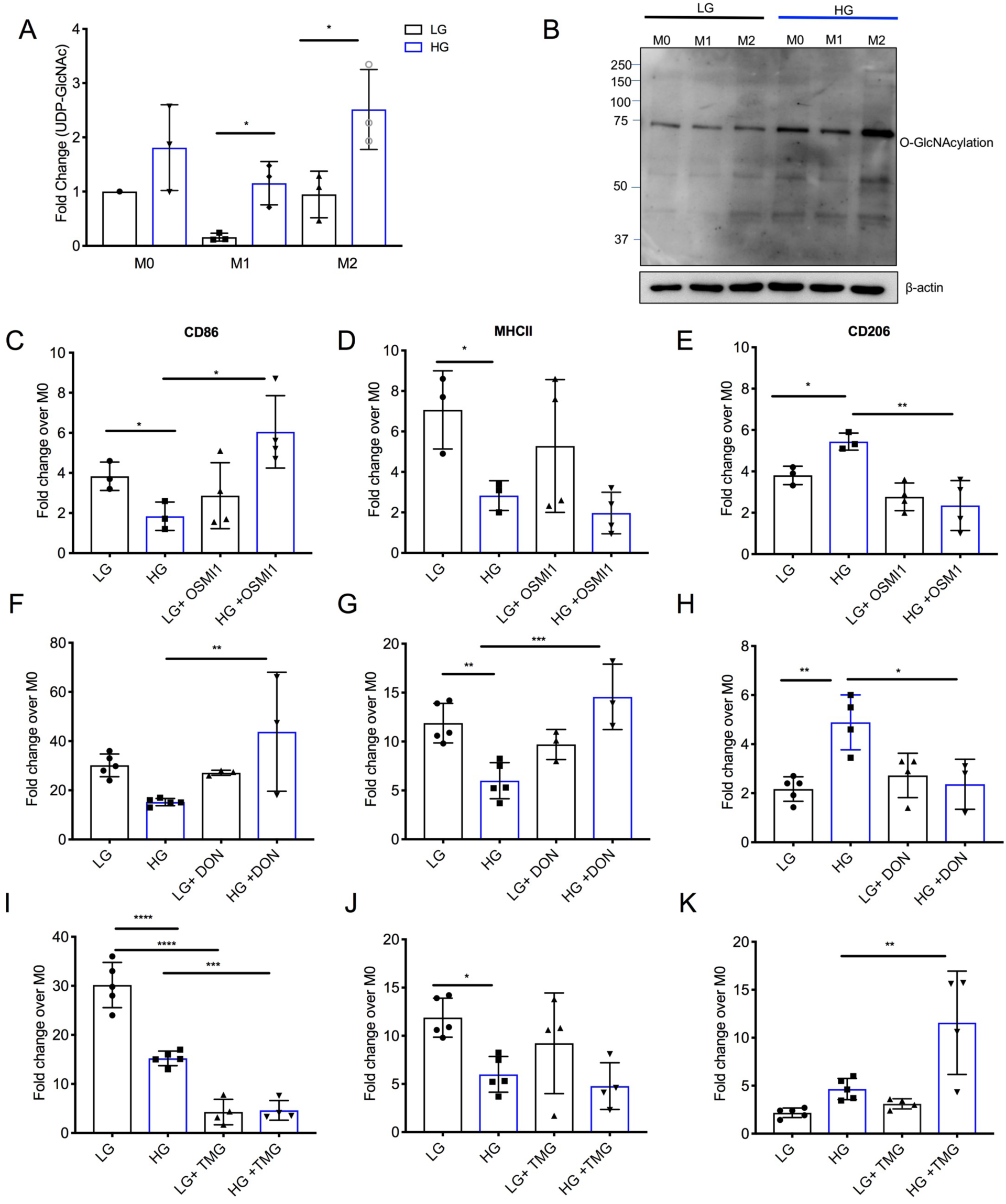
Flux through HBP induces M2 polarization. Bone marrow derived macrophages (BMDM) were differentiated with M-CSF in DMEM HG (high glucose, 25 mM) or LG (low glucose, 5mM. Macrophages were polarized to M1 with IFN-γ and LPS or to M2 with IL-4 during 48 hours. (A) The metabolites were extracted and the UDP-GlcNAc evaluated by HPLC and (B) the levels of O-GlcNAcylation by Western blotting. After the differentiation, BMDM were treated with OSMI1 (C–E), TMG (F-H) or DON (I–K) during 24 hours. Macrophages were polarized to M1 or to M2 during 48 hours. Macrophage polarization markers were analyzed by FACS. M1, CD86^+^ (C, F and I), MHC-II^+^ (D, G and J) and M2 CD206^+^ (E, H and K). Results are expressed as mean ± S.D. p values were calculated using Anova one-way with Tukey post-test. * *P* < 0.05, ** *P* < 0.01, *** *P* < 0.001.

To determine whether O-GlcNAcylation would induce an M2 phenotype in BMDMs, we inhibited the enzyme O-GlcNAc-transferase (OGT) using the inhibitor OSMI-1 (Fig. 4C–E). Treatment decreased overall cell O-GlcNAcylation (Fig. S4H) and reversed the effects induced by HG, leading to an increase in CD86 and MHC-II expression (Fig. 4C, D) and a decrease in CD206 (Fig. 4E). Applying the glutamine antagonist 6-diazo-5-oxo-L-norleucin (DON), which inhibits the rate-limiting enzyme of HBP glutamine fructose-6-phosphate amidotransferase (GFAT) (Fig. 4 F–H) and leads to a decrease in O-GlcNAcylation (Fig. S4I), showed similar effects.

Conversely, increasing O-GlcNAcylation by inhibiting OGA with Thiamet G (Fig. S4J), augmented the expression of CD206 (Fig. 4K) and decreased CD86 in both high and low glucose cultured macrophages (Fig. 4I). These experiments suggest that high glucose concentrations enhance M2 polarization in murine bone marrow-derived macrophages by increasing the flux of glucose through the HBP, resulting in an increase of O-GlcNAcylation. To further investigate the influence of O-GlcNAcylation on hyperglycemia-promoted tumor growth and macrophage fate *in vivo*, we treated hyperglycemic and euglycemic mice with the inhibitor of the HBP 6-diazo-5-oxo-l-norleucin (DON) thereby inhibiting O-GlcNAcylation. Treatment impaired tumor growth in both hyperglycemic and euglycemic mice (Fig. 5A) and resulted in a similar tumor growth in both groups, abrogating the effect of hyperglycemia (Fig. 5A). We observed a clear reduction of polarization to a pro-tumorgenic phenotype of macrophages (Fig. 5B, S5A) and an increase in M1 macrophage markers (Fig. S5B, C) in the tumor microenvironment of hyperglycemic mice. Confirming this finding, opposite effects were seen when mice were treated with the inhibitor of the enzyme that cleaves O-GlcNAc (OGA) Thiamet G increasing O-GlcNAcylation in mice. We found an increase in pro-tumorigenic M2-like macrophages (Fig. 5B), while no differences were observed in the CD206^+^ and MHC-II^+^ double positive population (Fig. S5D). Together, these results show that the inhibition of O-GlcNAcylation reverts the effect of hyperglycemia on tumor growth and intratumoral macrophage polarization *in vivo*, reprograming macrophages towards an antitumor phenotype.

**Figure 5:**
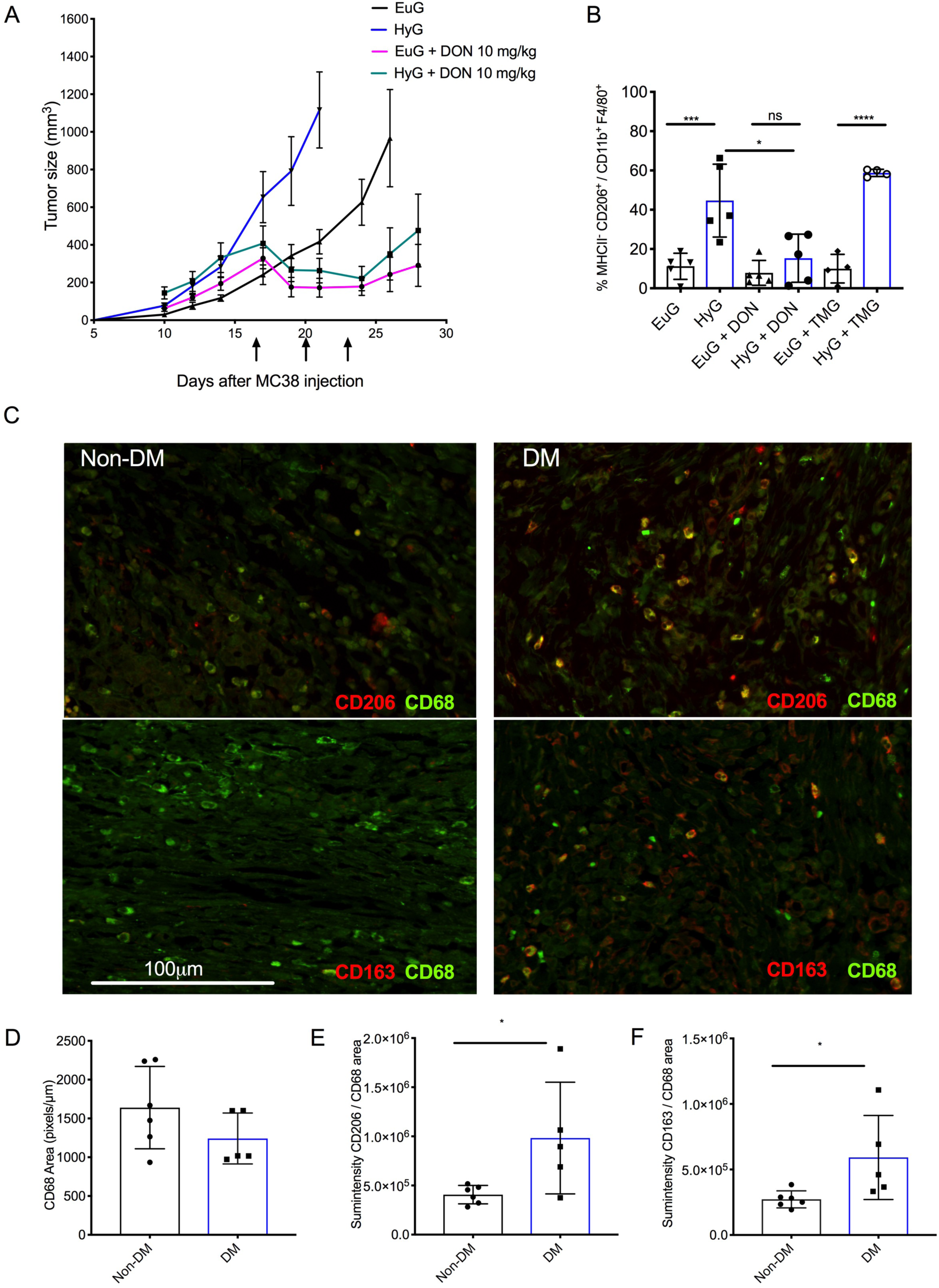
Flux through HBP induces M2 polarization. (A) Hyperglycemia (HyG) was induced in *vivo* by i.p. injection of streptozotocin (STZ) 150 mg/kg, while Euglyemic control (EuG) received the vehicle. When tumors reached 300-400 mm^3^ mice received DON (10 mg/mg) i.p. each 3 days (black arrows) and tumors were measured. (B) Macrophage polarization was analyzed after one single injection of DON (10 mg/kg) or TMG (20 mg/kg) when the tumors reached 300–400 mm^3^. Two days after the treatment, tumors were analyzed by cytometry using macrophage markers CD11b^+^F4/80^+^Gr1^−^ and polarization markers MHC-II and CD206 (n=4–5). (C) CRC samples from patients with diabetes mellitus type 2 (DM) or not (Non-DM) were stained for the macrophage marker CD68 (green) and M2 polarization markers CD206 and CD163 (red). The quantification was performed using the macrophage area, CD68 (D) and the sum intensity of CD206 (E), or CD163 (F) per CD68 area. Results are expressed as mean ± S.D. p values were calculated using Statistical Anova one-way with Tukey post-test. *** P < 0.001 and **** P < 0.0001 or unpaired Student’s t-test. * P < 0.05.

### Diabetes alters the tumor immune microenvironment in colorectal cancer

Hyperglycemia correlates with a worse prognosis in colorectal cancer [32] and our data from preclinical models provided evidence that hyperglycemia could induce intratumoral macrophage polarization to a protumorigenic M2-like phenotype. Analysis of TCGA gene expression data showed that the expression of M2 macrophage markers correlated with a reduced overall survival for all types of cancers (Fig. S5E) and that the increased expression of OGT was similarly associated with poor survival (Fig. S5F). To investigate if hyperglycemia was associated with a protumorigenic M2-like polarization of macrophages in colorectal cancer (CRC) patients, we analyzed samples from CRC patients with type 2 DM for changes in macrophage infiltration and polarization (Fig. 5C). No significant differences in the frequency of CD68^+^ intratumoral macrophages were observed between non-diabetic and diabetic patients (Fig. 5C). However, immunohistochemical analysis of CD68^+^ macrophages for the M2 markers CD206 (Fig. 5E and S6A) and CD163 (Fig. 5F and S6B) showed a higher frequency of M2-like macrophages in diabetic patients. This analysis suggests that hyperglycemic CRC patients to have a higher frequency of M2 macrophages, although larger studies are needed to further corroborate a direct association between DM2 and M2 macrophage polarization.

## Discussion

Here, we show that systemic and intratumoral hyperglycemia significantly alters the tumor immune microenvironment and increases the number of alternatively M2-like polarized tumor-promoting macrophages. We demonstrate that these macrophages enhance tumor growth by mediating immune evasion and secondary inhibition of the adaptive immune response. Intratumoral macrophages are known to support cancer progression by creating an inhibitory microenvironment and to strongly inhibit anti-tumor immunity [33–35]. M2-like polarized macrophages secrete mediators such as IL-10 and TGF-β leading to the dampening of adaptive anti-tumor immunity [33–35]. M2-like polarized TAMs are also defective of phagocytosing tumor cells [36–38]. Due to the heavy involvement in immune evasion and cancer progression, TAMs have become an interesting target for cancer immunotherapy. For example, blockade of CD47 on tumor cells or its receptor SIRPα on macrophages was successfully tested in preclinical models and antibodies against CD47 are currently in clinical development [36–38]. Targeting the recruitment and polarization of macrophages has also been tested by using blocking agents against CSF-1 or CSF-1R [39, 40]. In our model, treatment with anti-CSF1R led to a significant reduction in TAMs and M2-like polarization. This reduction of protumorigenic macrophages reversed the tumor-promoting effect of hyperglycemia, which clearly implicates intratumor macrophages in the observed effect.

Polarization of TAMs is not binary but rather a continuum [33–35], with LPS and IFNγ inducing an anti-tumoral M1 phenotype, while cytokines including IL-4 and IL-13 lead to an alternative, anti-inflammatory M2-like phenotype. Previous evidence showed that M2 polarization in murine macrophages is accompanied by an increase in UDP-GlcNAc [28]. We demonstrate that hyperglycemia increases M2 macrophage polarization through an increase in HBP flux and enhanced UDP-GlcNAc production. Inhibition of the HBP with DON, showed a reduction in M2 polarization and also abolished the effect of hyperglycemia *in vivo*, although the results of systemic DON treatment need to be interpreted cautiously due to its pleiotropic effects on many different cell types including direct effects on tumor cells. In further support of an important role of O-GlcNAcylation in hyperglycemia-driven M2 polarization, inhibition of the O-GlcNAc transferase by OSMI-1 inhibited the M2 polarization in hyperglycemic condition. O-GlcNAcylation has been associated in regulation of inflammatory responses and other immune reactions [21, 41].

Our data supports a role for O-GlcNAcylation in regulating macrophage polarization and differentiation, with increased O-GlcNAcylation directing macrophages towards M2-like phenotype. We show for the first time, that intratumoral hyperglycemia leads mediates cancer immune evasion by O-GlcNAc-mediated macrophage polarization. In agreement, recent work has shown that lipopolysaccharide (LPS) activated macrophages (M1 phenotype) display reduced HBP activity and protein O-GlcNAcylation [31]. Previous studies have reported how O-GlcNAc cycling can influence macrophage function. Depletion of OGT in the human macrophage cell line THP-1 induces M1 polarization, whereas M2 genes were down-regulated [42]. In addition, pharmacological inhibition of OGA in an experimental model of ischemic stroke showed an increased expression of M2 markers and decreased expression of M1 markers in microglia cells [43]. OGT-mediated O-GlcNAcylation of STAT3 in macrophages inhibits STAT3 phosphorylation and IL-10 production [44]. Several proteins which regulate macrophage function, i.e. Akt/mTOR, HIFα, as well as the NFκB and NFAT families [35, 45–47], were shown to be *O*-GlcNAcylated in other models. Together these studies point to OGT as an important mediator of M2 polarization in macrophages.

A limitation of our study is the use of STZ to induce hyperglycemia, since the model induces hypoinsulinemia, where the acute type of hyperglycemia resembles more the situation in patients with type 1 DM. Future experiments will make use of other DM models, including diet-induced mouse models that mimic type 2 DM. We have also not identified exact target proteins of O-GlcNAcylation in our experiments, but have focused on overall levels of O-GlcNAcylation. Further studies are needed to determine the distinct influence of enhanced O-GlcNAcylation on different signaling pathways.

Our results support a metabolic reprogramming of the tumor immune microenvironment by hyperglycemia resulting in an immune-inhibitory, tumor-promoting condition mediated by increased M2-like macrophage polarization. Accordingly, diabetic patients with CRC had an increased number of M2-like polarized macrophages compared to CRC patients without diabetes. While our study advocates for a strict control of glycemia in diabetic cancer patients, further functional analyses and clinical examinations are needed to determine the target range of blood glucose in cancer patients. One could also speculate that diabetic cancer patients could benefit from an additional macrophage-targeting agent such as CSF-1R blockade. Furthermore, the description of O-GlcNAcylation mediating hyperglycemia-induced tumor progression provides new targets for cancer immunotherapy, including enzymes involved in the HBP and O-GlcNAcylation. We demonstrated that OGT inhibition reprograms TAMs to an antitumor profile. By directly targeting macrophages, the specific delivery of OGT inhibitors could be achieved, greatly reducing the toxicity of the compound and influencing the tumor microenvironment only, reducing potential side effects.

## Acknowledgments

This work was supported by funding from the Goldschmidt-Jacobson Foundation (to H.L.), the Promedica Foundation (to M.A.S. and A.Z.), Krebsliga Beider Basel (KLBB, to H.L.), Schoenemakers Foundation (to H.L.), Swiss Government Excellence Scholarships for Foreign Scholars and Artists (FCS, to N.R.M.), Coordenação de Aperfeiçoamento de Pessoal de Nível Superior (CAPES to N.R.M. and R.C.S.), Conselho Nacional de Desenvolvimento Científico e Tecnológico (CNPq to A.R.T, I.A.O., N.R.M. and W.B.D.), Fundação de Amparo à Pesquisa do Estado do Rio de Janeiro (FAPERJ to A.R.T.) and Fundação do Câncer (to W.B.D.). We thank the Centro de Espectrometria de Massas de Biomoléculas (CEMBIO) and plataforma de Imuno-análise (PIA) (UFRJ, Rio de Janeiro, Brazil). We also thank all the patients that allowed to use their material and made this work possible.

## Author contributions

N.R.M., A.R.T., and H.L. designed the project. N.R.M., M.A.S., W.B.D., A.Z., A.R.T., and H.L. planned the experiments. A.F., R.C.S. and M.T. processed CRC sample, C.S.B analyzed CRC biopsies, N.R.M., A.F., M.A.S., I.O., N.K., A.S., and R.F. performed experiments. N.R.M. and M.A.S. analyzed the results. N.R.M., A.R.T., and H.L. wrote the manuscript. All authors read the manuscript and gave comments.

