## Supplemental Data for "Hyperglycemia enhances cancer immune evasion by inducing alternative macrophage polarization through increased O-GlcNAcylation"

### Material and Methods

#### Antibodies

The following antibodies were used for flow cytometry staining of murine cell: Fc-block antibody anti-CD16/36 (clone 2.4G2), anti-CD45 PercPcy5 (clone 30-F11), anti-Gr1 PEcy7 (clone RB6-8C5), anti-CD11b FITC/BV605 (clone M1/70), anti-F4/80 BV421 (clone BM8), anti-MHCII APCcy7 (clone M5/114.15.2), anti-CD3-PE594 (clone 17A2), anti-CD4 BV605 (clone RM4-5), anti-FoxP3 APC (clone 3G3), anti-CD25 PE (clone 3C7), anti-CD19 APCcy7 (clone 6D5), anti-CD11c APC (clone HL3), anti-Ly6C BV605 (clone AL-21), anti-Ly6G PEcy7 (clone 1A8), anti-CD86 APC (clone GL-1) anti-CD206 PE (clone C068C2). All the antibodies for flow cytometry were purchased from BD Biosciences or Biolegend.

#### Sialylation deficient MC38 cells

MC38 GNE (UDP-N-acetylglucosamine 2-epimerase/N-acetylmannosamine kinase) deficient cells were generated by CRISPR/CAS9 technique as previously described [1]. Briefly, guide RNAs were designed with e-crisp.org and synthesized by Microsynth AG. Guide RNAs were cloned into the pX458 vector (Addgene). A transient transfection of MC38 cells with subsequent single-cell sorting and screening for cell-surface sialylation were performed. Five MC38 GNE-deficient clones were pooled to avoid clonal selection.

#### Siglec E-Fc production

The Siglec-Fcs were produced a previously described [2]. Briefly, transfected 293T HEK cells were cultured in serum free medium and the supernatant collected after 72 hours. Siglec-Fc was isolated from supernatants with protein A.

#### Siglec E KO mice

The Siglec-E deficient mouse (EKO mouse) was received from Dr.Varki (UCSD) previously described. The mice were bred and backcrossed in-house to our local C57BL/6 strain, in heterozygous breedings for more than 9 generations. Animals were housed under specific pathogen-free conditions.

#### Survival analysis with TCGA data

Harmonized RNA-Seq data from The Cancer Genome Atlas (TCGA) database were retrieved using the software TCGAbiolinks [3]. The normalization method chosen for the expression values was the FPKM-UQ. The survival analysis was performed on the patients of all cancers which were split in two categories, top and low 50% of the list of patients ranked according to expression of the OGT gene. The p-value for Kaplan-Meier curves was calculated with the log rank test. The proportions of the M1 and M2 macrophages infiltrated in the tumors of the TCGA database were retrieved with the CIBERSORT method [4]. Survival analysis was performed as described above but using either the M1 or M2 frequency to split the patients.

**Nanostring analysis**

RNA extraction was performed of tumor embedded in OCT. 5-10 10 µM sections of tumor tissue were lysed in 600 µl RLT buffer using the TissueLyserLT (Qiagen). RNA was extracted using the RNeasy Mini extraction Kit (Qiagen) followed by nCounter Low RNA Input Amplification Protocol (nanoString). The Nanostring data was normalized using the housekeeping genes selected by the geNorm [5] algorithm from the Advanced Analysis of the nSolver software. The limma package was used to perform the differential expression analysis between the normoglycemic and hyperglycemic mice. The data were log2-transformed with the voom function from limma in order to take into account the mean-variance relationship [6]. The volcano plot shows in red the genes with a p-value $< 0.05$. The labelled genes are the ones with a p-value $< 0.05$ and absolute logFC $>0.75$.

### Supplementary Figures

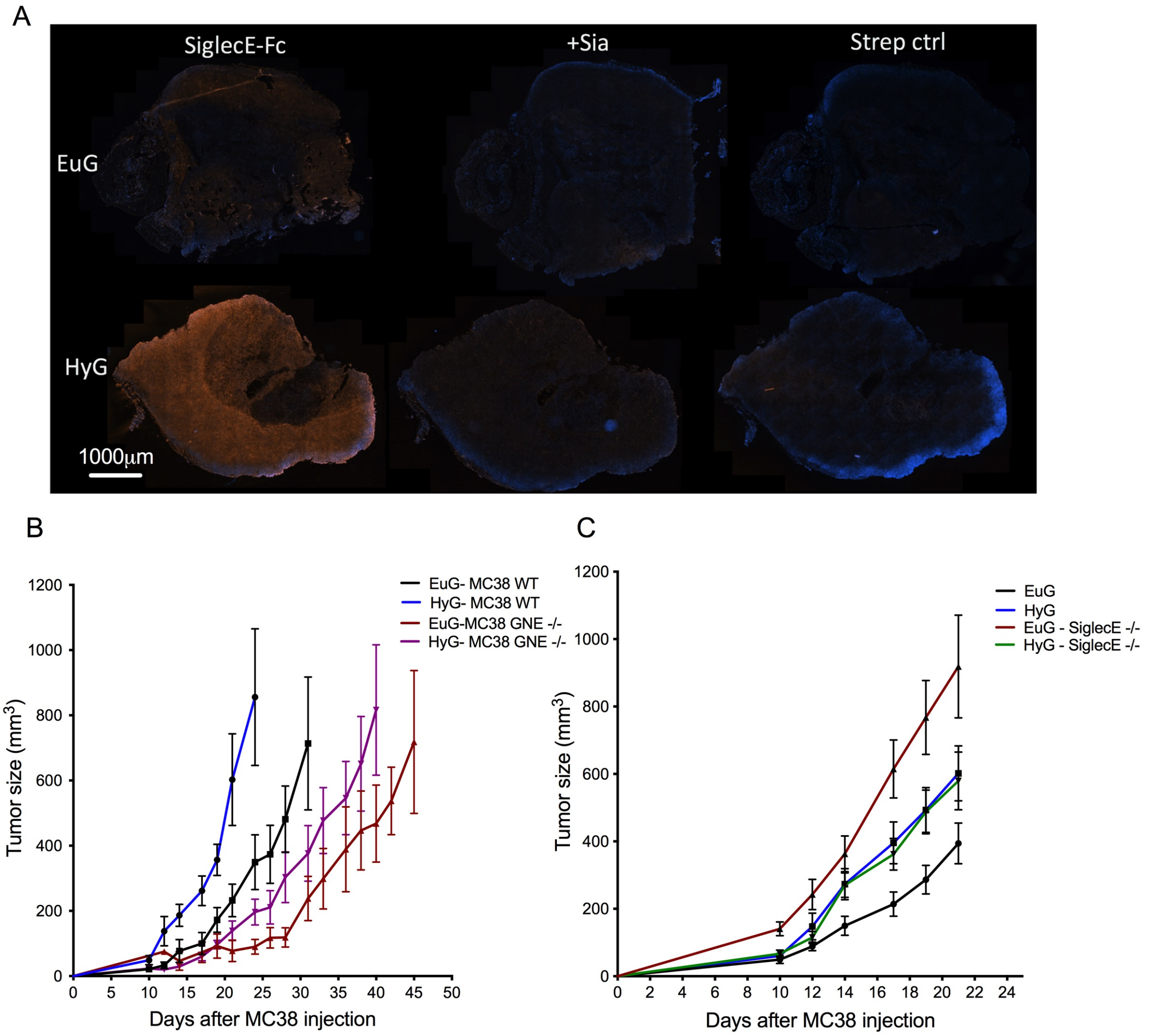


**Figure S1- Sialylation is not a key mechanism in hyperglycemia-induced tumor growth.** Tumor cells were injected one week after an i.p injection of STZ 150 mg/kg, hyperglycemic mice (HyG) or vehicle Euglyemic group (EuG). (A) The implanted tumors from EuG or HyG mice were sectioned and stained for SiglecE recombinant protein fusionated with a Fc moiety, accessing the Siglec E ligands in the tumor by scanning the whole tumor tissue, the controls were made treating the tumor with sialidase (sia) or using only streptavidin as secondary. (B) Tumor growth curves comparing MC38 wild-type (WT) and MC38 GNE -/-, knocked out to GNE (UDP-N-acetylglucosamine 2-epimerase/N-acetylmannosamine kinase), treated with STZ (HyG-MC38 WT and HyG-MC38 GNE -/-) or non-treated (EuG- MC38- WT and EuG-MC38 GNE-/-). (C) Growth curve using C57bl/6 mice knocked out to SiglecE, treated with STZ (HyG-SiglecE -/-, n=8) or non-treated (EuG-SiglecE-/-, n=9) and the control C57bl/6 mice treated with STZ (HyG) and non-treated (EuG).

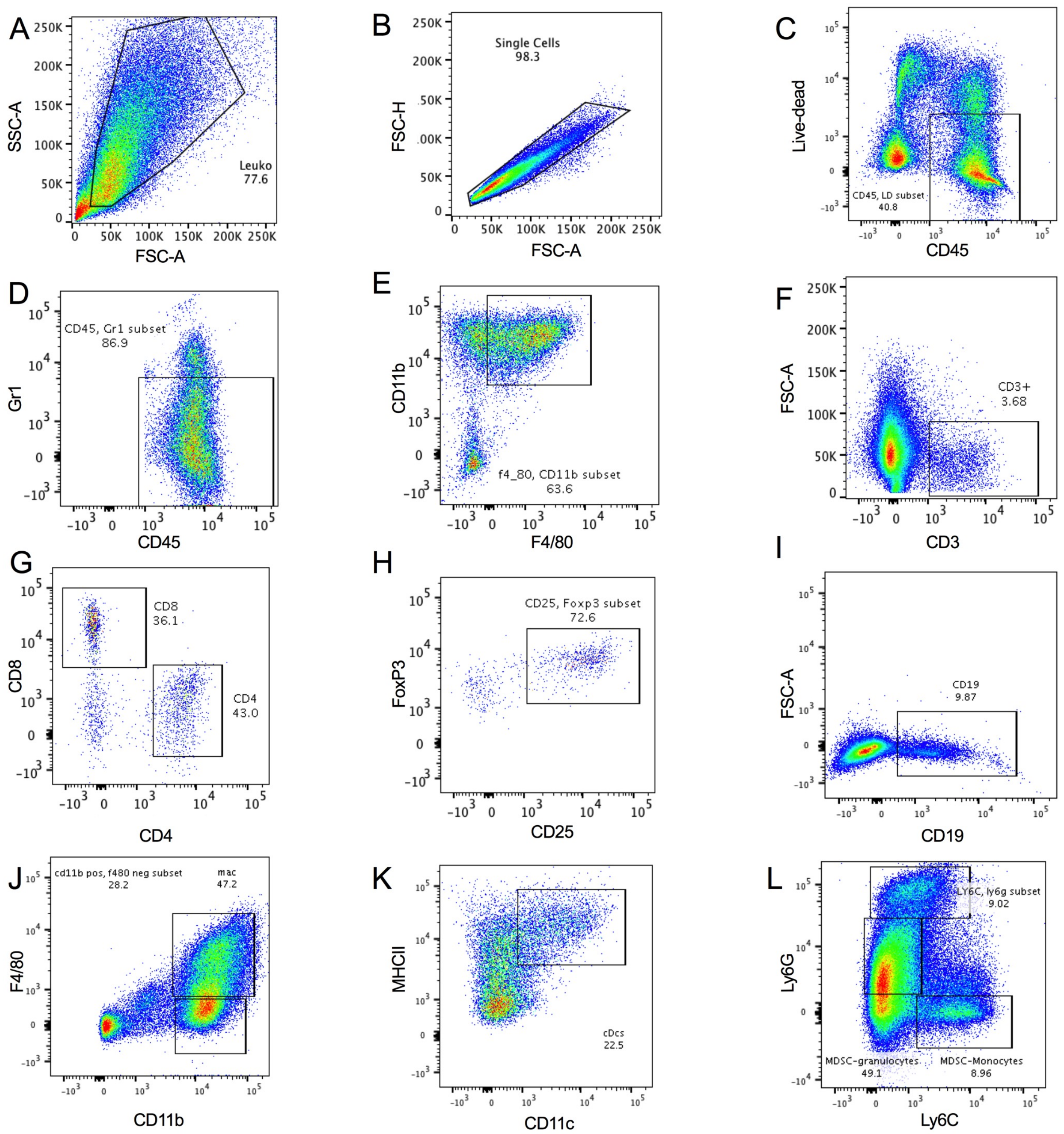


**Figure S2- Gating strategy to determine tumor cell infiltrates.** All the tumor inflammatory cells were gated on Side Scatter and Forward Scatter (A), singlets (B) and alive CD45+ cells (C). The tumor associated macrophages were also pre-gated on Gr1 negative (D) and $CD11b^{+}$ $F4/80^{+}$ cells (E). The tumor infiltrated lymphocytes are all CD3+ cells (F), gated on $CD8^{+}$ and $CD4^{+}$ (G). The T regulatory cells were analysed using the $CD4^{+}$ cells to quantify $CD25^{+}$ and $FoxP3^{+}$. B cells were gated on $CD19^{+}$ cells (I). Dendritic cells were pre-gated out of F4/80 population and kept the CD11b+ cells (J) and then $MHCII^{+}$ and $CD11c^{+}$(K). The Myeloid-derived suppressor cells inside CD11b+ cells and gates in three different gates $Ly6C^{lo}Ly6G^{hi}$, $Ly6C^{lo}Ly6G^{int}$ and $Ly6C^{hi}Ly6G^{lo}$ (L).

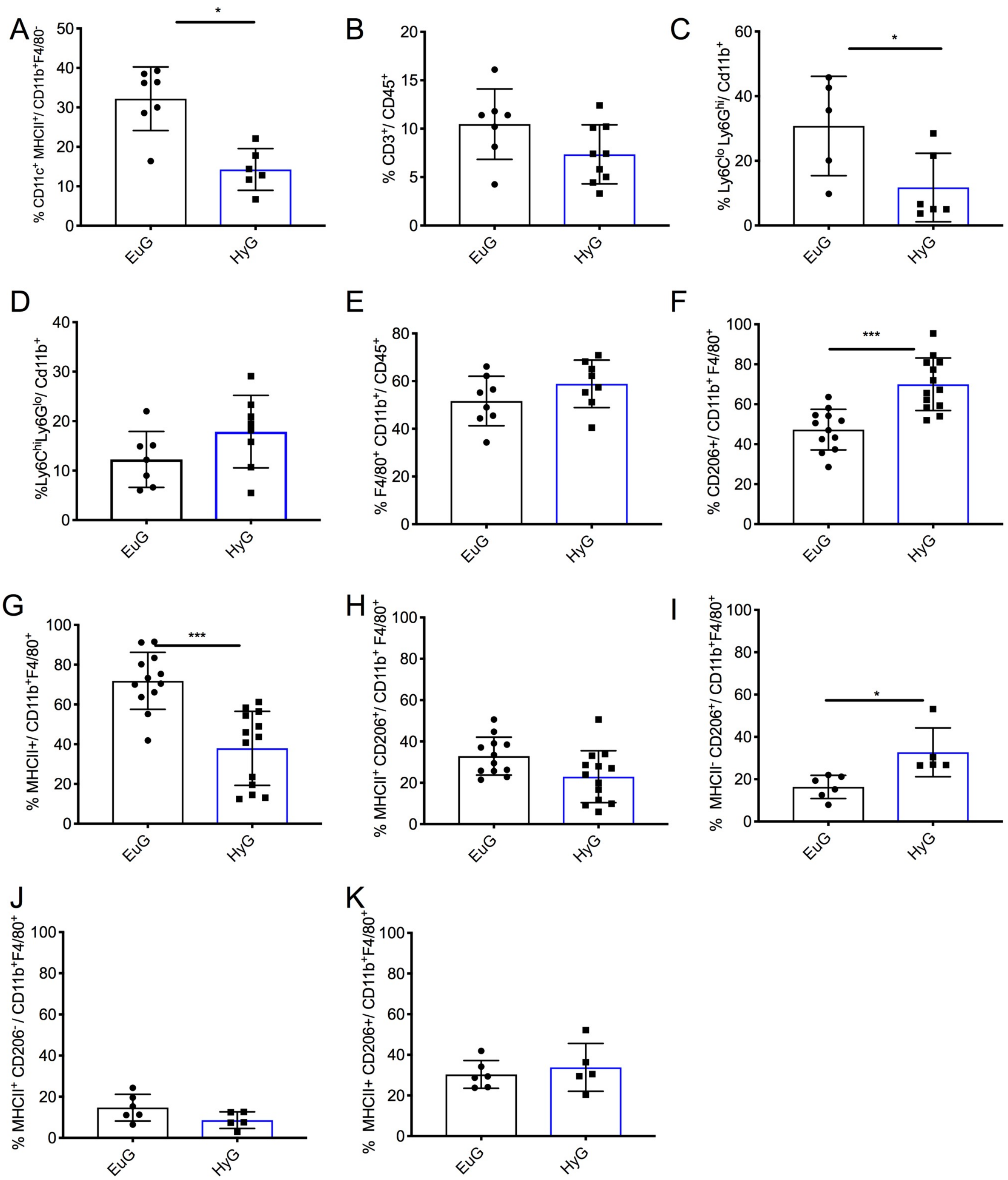


**Figure S3- Hyperglycemia affects anti-tumoral immunity.** Cell infiltrates were determined in MC38 and B16D5 solid tumors. (A) Dendritic cells (CD11c$^+$MHCII$^+$) were previously gated on CD11b+ F4/80$^-$ (n=8-6). Frequency of tumor infiltrated (B) lymphocytes CD3$^+$ (n=7-9) in MC38 tumors. The Myeloid-derived suppressed cells were performed using CD11b+ cells, divided in two different populations (C) Ly6C$^{lo}$Ly6G$^{hi}$ and (D) Ly6C$^{lo}$Ly6G$^{hi}$ (n=7-9). (E) The MC38 tumor associated macrophages (n=12-13) were analysed to the populations (F) CD206+, (G) MHCII+ and (H) MHCII+ CD206+. The B16D5 tumor associated macrophages (CD45+F4/80+CD11b+Gr1-) were analysed according the markers MHCII and CD206 (n=6-5), gates on (I) $MHCII^{-}CD206^{+}$, (J) $MHCII^{+}CD206^{-}$ and (K) $MHCII^{+}CD206^{+}$. Statistical analysis by unpaired Student's t- test. * $P<0.05$ and *** $P< 0.001$.

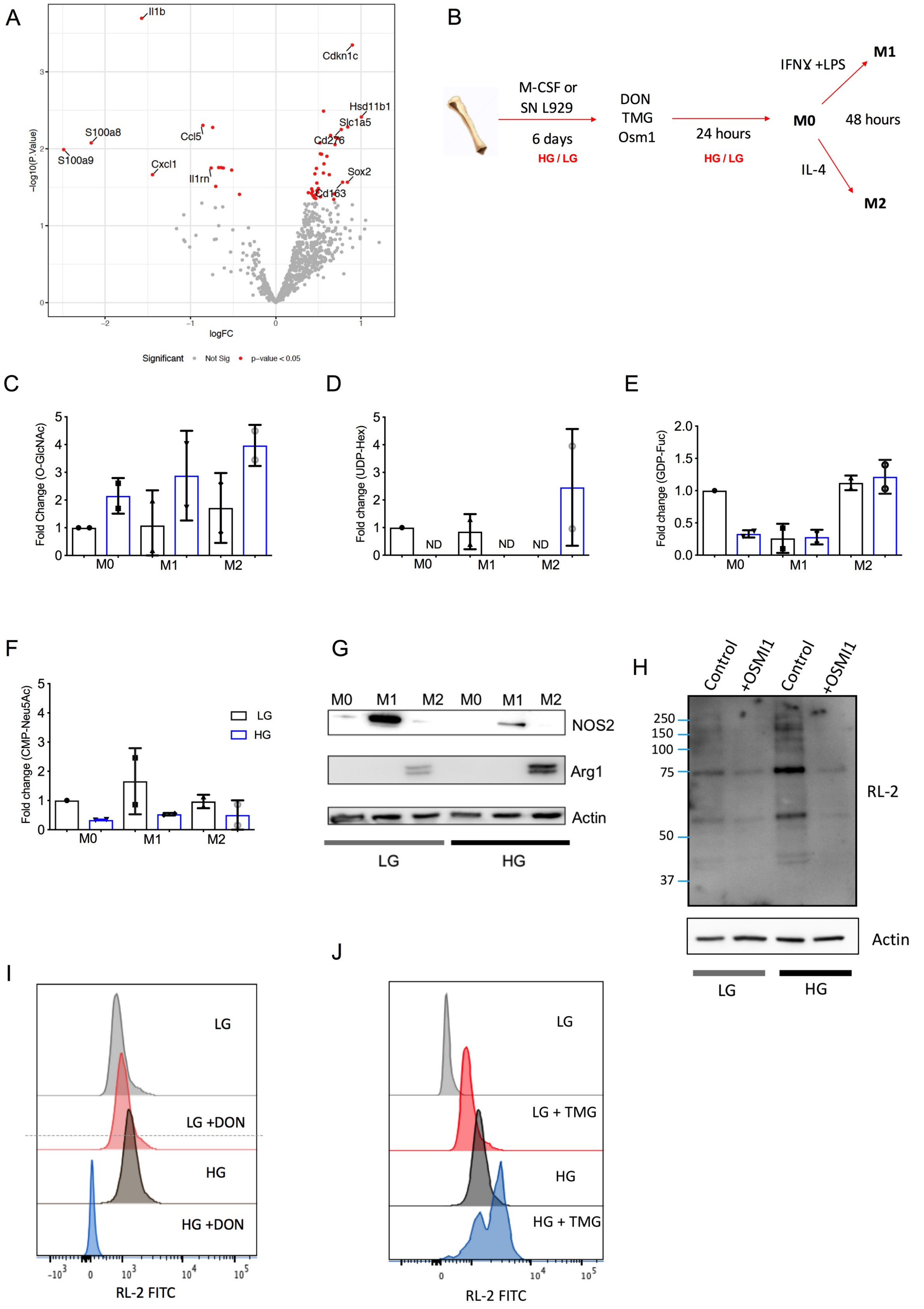


**Figure S4- Hyperglycemia affects macrophage polarization through O-GlcNAcylation.** Volcano plot of differentially expressed genes between Euglyemic and Hyperglicemic MC38

tumors (n=3-4). (A) RNA was extracted from MC38 tumors and analyzed by Nanostring platform. The volcano plot shows in red the genes with a p-value < 0.05. The labelled genes are the ones with a p-value < 0.05 and absolute logFC >0.75. (B) BMDM differentiated with L929 supernatant or M-CSF were polarized in DMEM HG (high glucose, 25 mM) or LG (low glucose, 5mM). (C) The quantification of O-GlcNAcylation using the antibody RL-2 by western blotting and the metabolites UDP-Hexoses (D), GDP-Fucose (E) and CMP-Neu5Ac (F) were extracted and analysed measured by mass spectrometry. (G) The levels of Arginase1 (Arg1) and Nitric oxide synthase 2 (NOS2) were analysed in the polarized macrophages M1 and M2. O-GlcNAcylation was accessed after 24 hours treatment of (H) OSMI1, (I) DON and TMG (J).

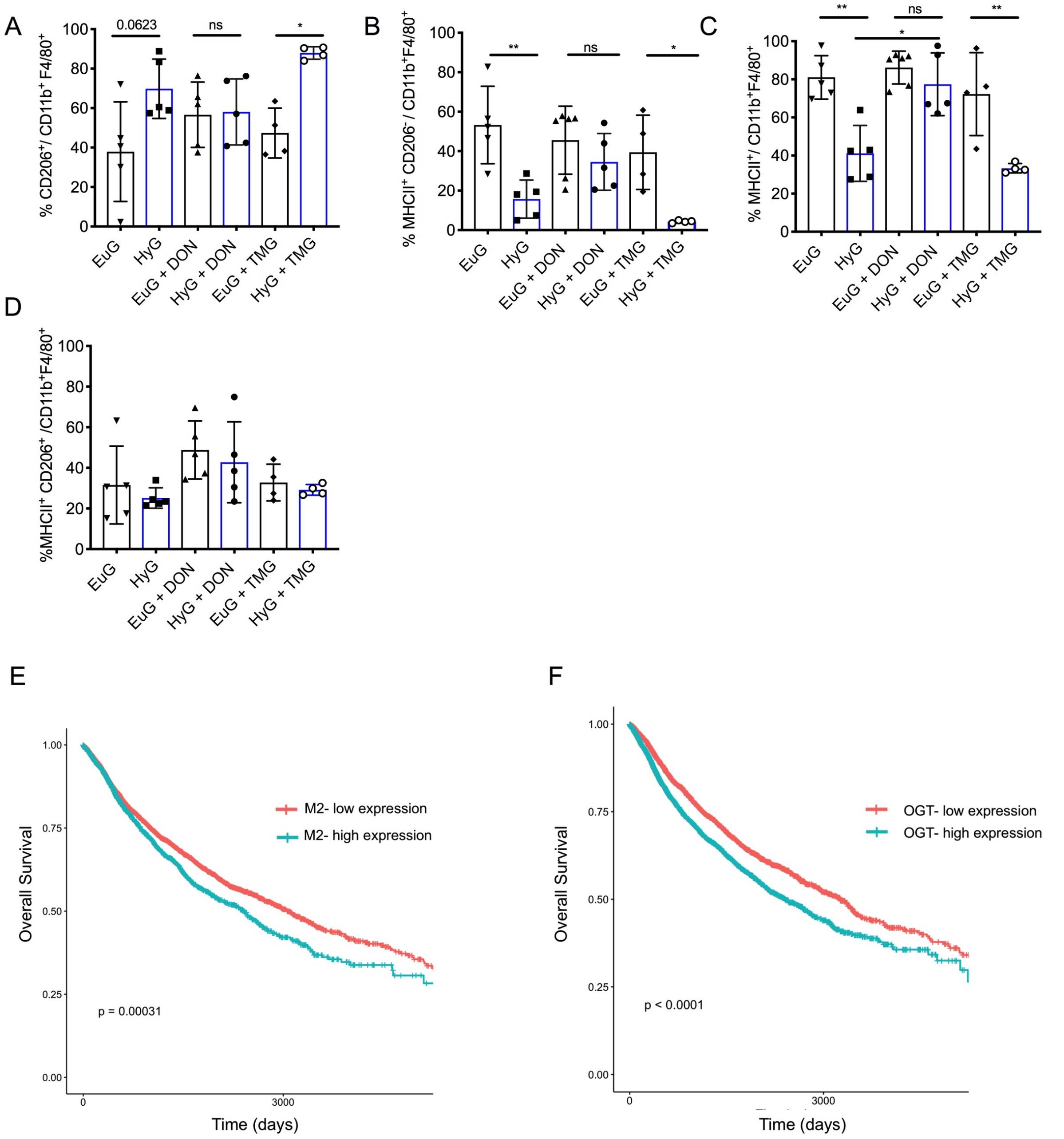


**Figure S5 - Flux through HBP induces M2 polarization.** Hyperglycemia (HyG) was induced in *vivo* by i.p. injection of streptozotocin (STZ) 150 mg/kg, while Euglyemic control (EuG) received the vehicle. When tumors reached 300-400 mm$^3$ mice received DON (10 mg/mg) i.p. each 3 days and tumors were measured. (B-D) Macrophage polarization was analysed after one single injection of DON (10 mg/kg) or TMG (20 mg/kg) when the tumors reached 300-400 mm$^3$. Two days after the treatment, tumors were analyzed by cytometry using macrophage markers CD11b$^+$F4/80$^+$Gr1$^-$ and polarization markers MHCII and CD206 (n=4-5). Results are expressed as mean ± S.D. p values were calculated using Statistical Anova one-way with Tukey post-test. * P < 0.05, ** P < 0.01. Harmonized RNA-Seq data from The Cancer Genome Atlas (TCGA) database were retrieved using the software

TCGAbiolinks. The survival analysis was performed on the patients of all cancers which were split in two categories, top and low 50% of the list of patients ranked according to expression of the (E) M2 or (F) OGT genes.

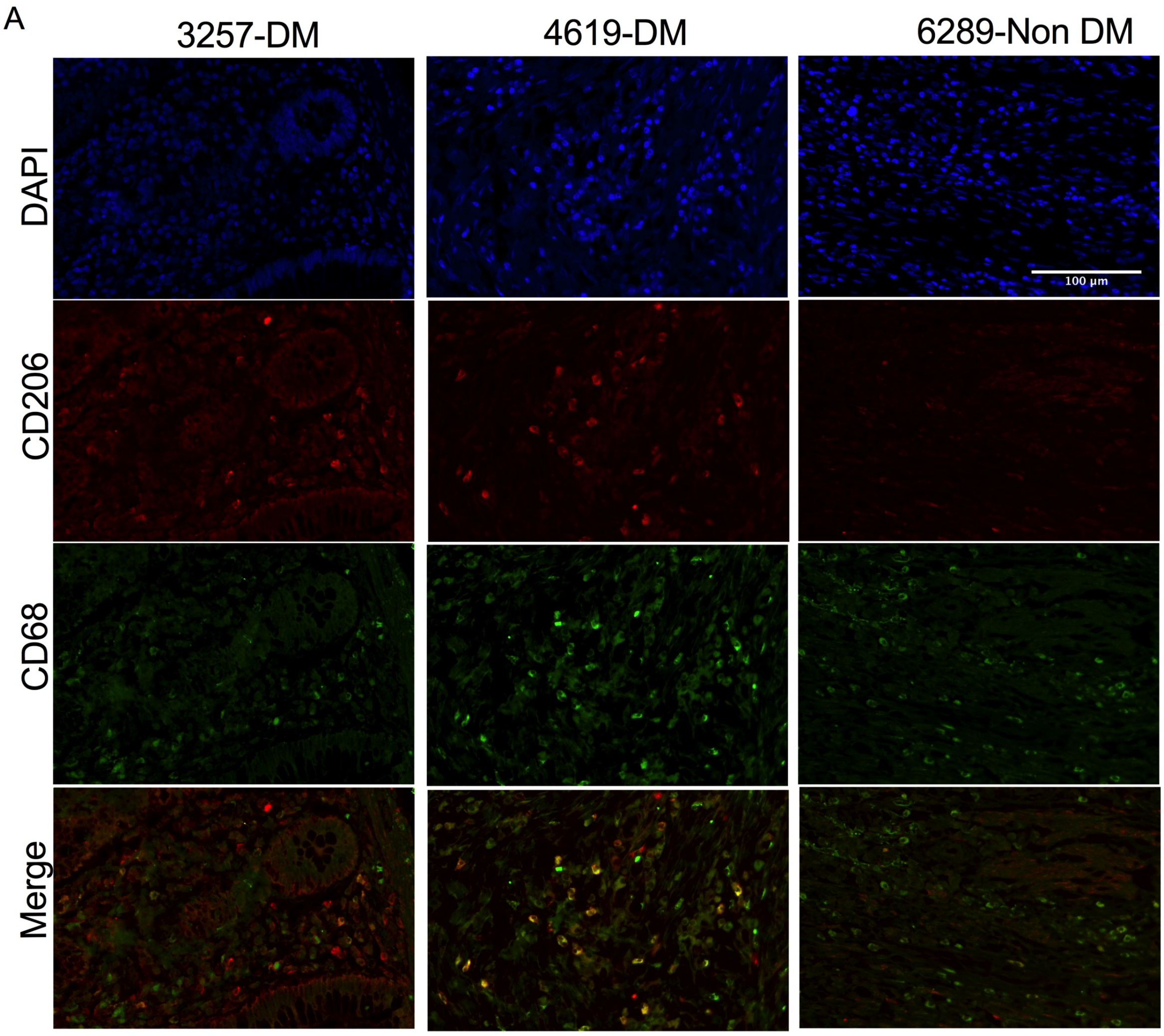
A
3257-DM
4619-DM
6289-Non DM
DAPI
CD206
CD68
Merge
100 µm

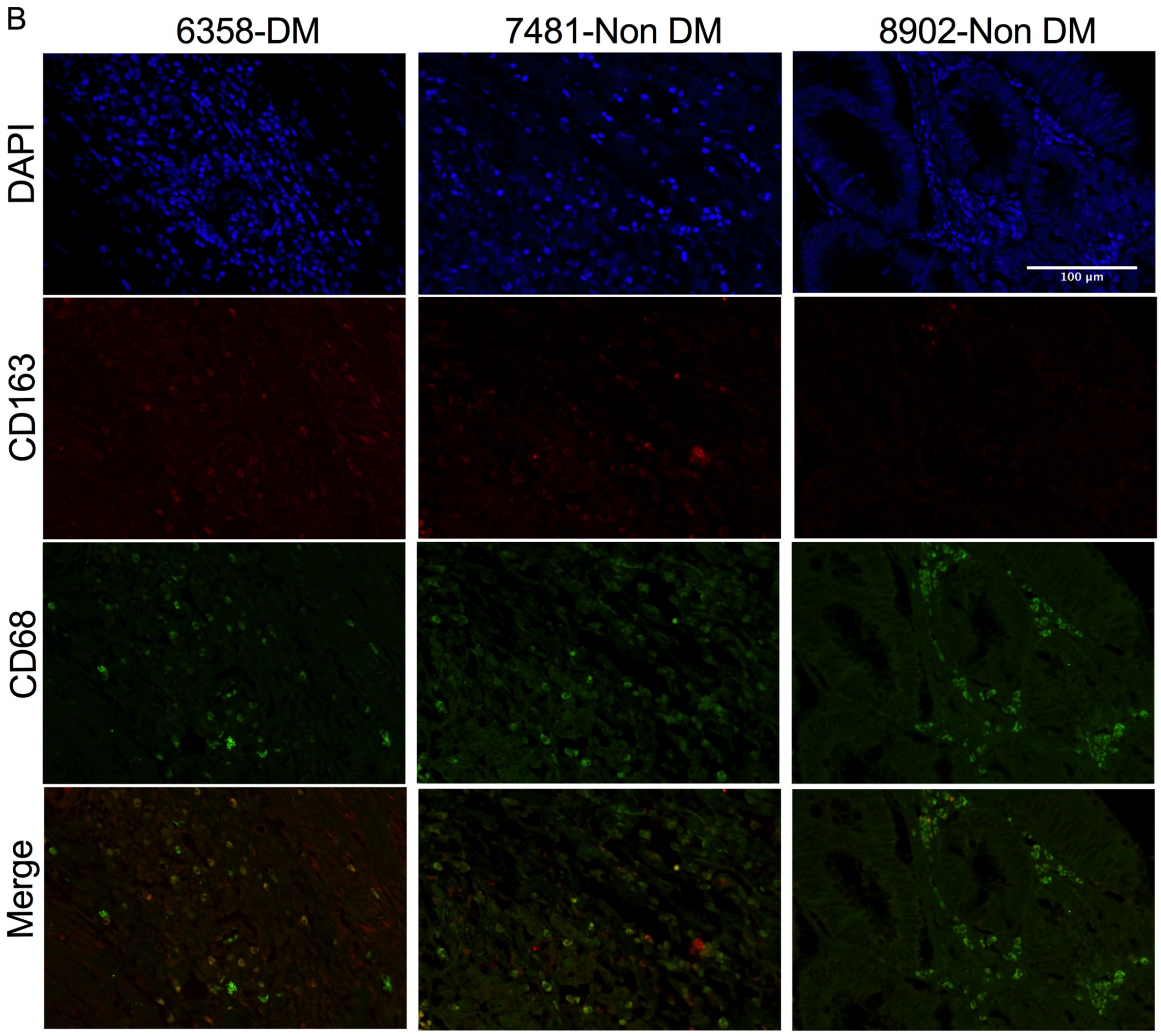


**Figure S6 - Diabetes induces M2 polarization in colorectal cancer**. CRC samples from patients with diabetes mellitus type 2 (DM) or not (control) were stained for the macrophage marker CD68 (green) and M2 polarization markers (A) CD206 and (B) CD163 (red). The nuclei were stained using mounting medium with DAPI (blue).
